# Giotto, a toolbox for integrative analysis and visualization of spatial expression data

**DOI:** 10.1101/701680

**Authors:** Ruben Dries, Qian Zhu, Rui Dong, Chee-Huat Linus Eng, Huipeng Li, Kan Liu, Yuntian Fu, Tianxiao Zhao, Arpan Sarkar, Feng Bao, Rani E George, Nico Pierson, Long Cai, Guo-Cheng Yuan

## Abstract

The rapid development of novel spatial transcriptomic and proteomic technologies has provided new opportunities to investigate the interactions between cells and their native microenvironment. However, effective use of such technologies requires the development of innovative computational tools that are easily accessible and intuitive to use. Here we present Giotto, a comprehensive, flexible, robust, and open-source toolbox for spatial transcriptomic and proteomic data analysis and visualization. The data analysis module provides end-to-end analysis by implementing a wide range of algorithms for characterizing cell-type distribution, spatially coherent gene expression patterns, and interactions between each cell and its surrounding neighbors. Furthermore, Giotto can also be used in conjunction with external single-cell RNAseq data to infer the spatial enrichment of cell types from data that do not have single-cell resolution. The data visualization module allows users to interactively visualize the gene expression data, analysis outputs, and additional imaging features, thereby providing a user-friendly workspace to explore multiple modalities of information for biological investigation. These two modules can be used iteratively for refined analysis and hypothesis development. We applied Giotto to a wide range of public datasets encompassing diverse technologies and platforms, thereby demonstrating its general applicability for spatial transcriptomic and proteomic data analysis and visualization.

## Introduction

Most tissues consist of multiple cell types that operate together to perform their functions. The behavior of each cell is in turn mediated by its tissue environment. With the rapid development of single-cell RNAseq (scRNAseq) technologies in the last decade, most attention has gone to unravelling the composition of cell types with each tissue. However, recent studies have also shown that identical cell types may have tissue-specific expression patterns ^1,2^, indicating that the tissue environment plays an important role in mediating cell states. Since spatial information is lost during the process of tissue dissociation and cell isolation, the scRNAseq technology is intrinsically limited for studying the structural organization of a complex tissue and interactions between cells and their tissue environment.

Recently, a number of technological advances have enabled transcriptomic/proteomic profiling in a spatially resolved manner ^3–14^ such that cellular features (for example transcripts or proteins) can be assigned to single cells for which the original cell location information is retained (**Fig. 1A, inset**). Applications of these technologies have revealed distinct spatial patterns that previously are only inferred through indirect means ^15,16^. There is an urgent need for standardized spatial analysis tools that can facilitate comprehensive exploration of the current and upcoming spatial datasets ^17,18^. To fill this important gap, we present the first comprehensive, standardized and user-friendly toolbox, called Giotto, that allows researchers to process, (re-)analyze and interactively visualize spatial transcriptomic and proteomic datasets. Giotto implements a rich set of algorithms to enable robust spatial data analysis, and further provides an easy-to-use workspace for interactive data visualization and exploration. As such, the Giotto toolbox serves as a convenient entry point for spatial transcriptomic/proteomic data analysis and visualization. We have applied Giotto to a wide range of public datasets to demonstrate its general applicability.

**Figure 1.**
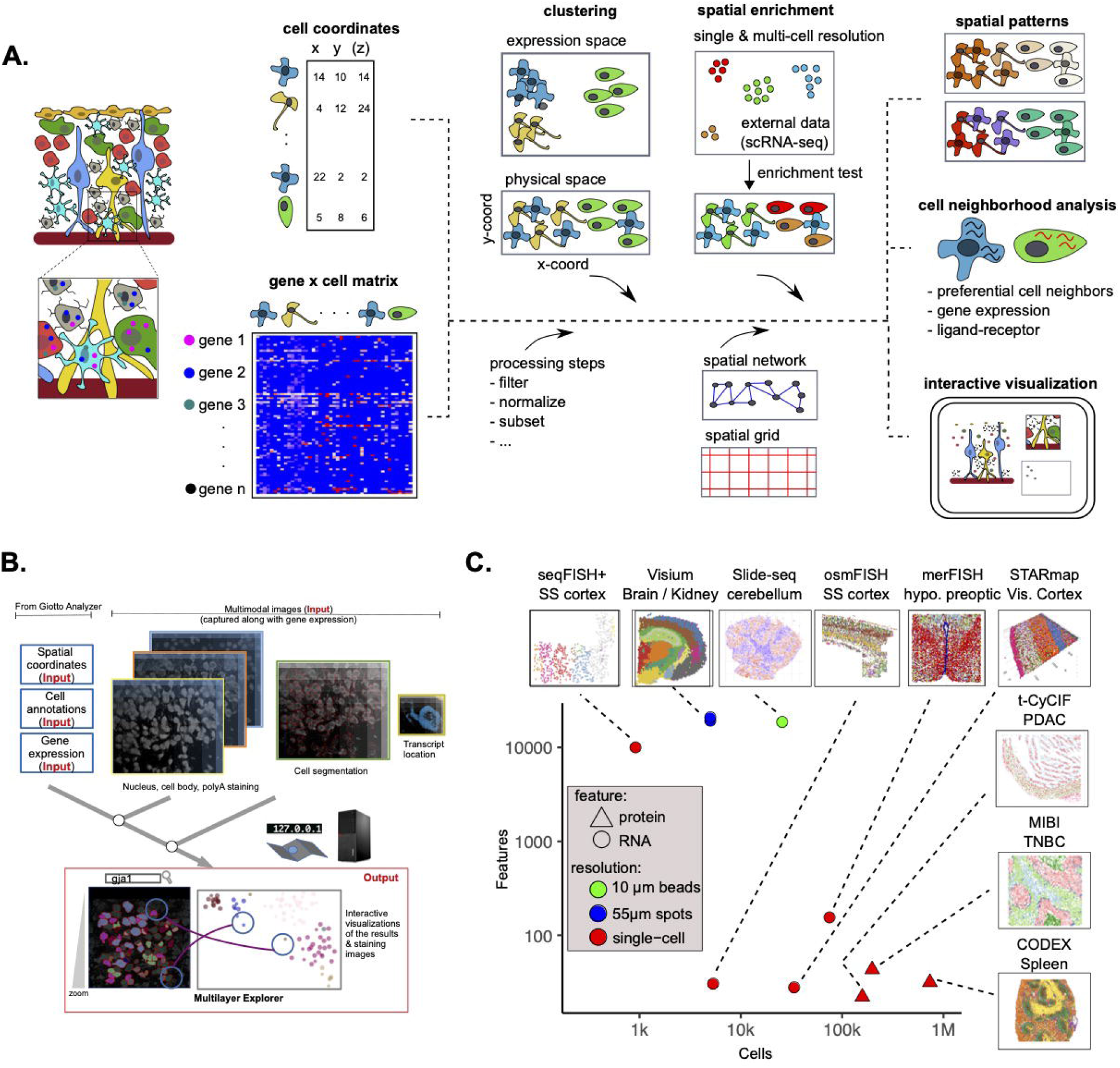
The Giotto framework to analyze and visualize spatial expression data. **A**. Schematic representation of the Giotto workflow to analyze and visualize spatial expression data. Giotto Analyzer requires a count matrix and physical coordinates for the corresponding cells. It follows standard scRNA-seq processing and analysis steps to identify differentially expressed genes and cell types. In the following step a spatial grid and neighborhood network is created which is further used to incorporate the spatial information of the single-cell environment and which is used for spatial analysis. **B**. Cell coordinates, annotations, and clustering information are utilized and incorporated in the Giotto Viewer. This interactive viewer allows users to explore the link between cells’ physical positions and their clustering pattern in the expression space (UMAP or tSNE). The addition of raw subcellular transcript coordinates, staining images or cell segmentation information is also supported. **C**. Overview of the selected broad range of different spatial technologies and datasets which were analyzed and visualized with Giotto. For each dataset the number of features (genes or proteins) and number of cells are shown before filtering. The technologies depicted are sequential fluorescence *in situ* hybridization plus (seqFISH+), Visium 10X (Visium), Slide-seq, cyclic-ouroboros single-molecule fluorescence *in situ* hybridization (osmFISH), multiplexed error-robust fluorescent *in situ* hybridization (merFISH), spatially-resolved transcript amplicon readout mapping (STARmap), tissue-based cyclic immunofluorescence (t-CyCIF), Multiplex Ion Beam Imaging (MIBI), and CO-Detection by indexing (CODEX).

## Results

### Overview of the Giotto toolbox

Giotto provides a comprehensive spatial analysis toolbox that contains two independent yet fully integrated modules **(Fig. 1A, B**). The first module (Giotto Analyzer) provides step-by-step instructions about the different steps in analyzing spatial single-cell expression data, whereas the second module (Giotto Viewer) provides a responsive and interactive viewer of such data on the user’s local computer. These two modules can be used either independently or iteratively.

Giotto Analyzer requires as minimal input a gene-by-cell count matrix and the spatial coordinates for the centroid position of each cell (**Fig. 1A**). At the basic level, Giotto Analyzer can be used to perform common steps often similar to scRNAseq analysis, such as pre-processing, feature selection, dimension reduction and unsupervised clustering; on the other hand, the main strength comes from its ability to integrate gene expression and spatial information in order to gain insights into the structural and functional organization of a tissue and its expression patterns. To this end, Giotto Analyzer creates a spatial grid and neighborhood network connecting cells that are physically close to each other. These objects function as the basis to perform analyses that are associated with cell neighborhoods.

Giotto Analyzer is written in the popular language R. The core data structure is the giotto object, which is specifically designed for spatial expression data analysis based on the flexible S4 object system in R. The giotto object stores all necessary (spatial) information and is sufficient to perform all calculations and analyses (**Supplementary Fig. 1A**). This allows the user to quickly evaluate and create their own flexible pipeline for both spatial visualization and data analysis. The Giotto Viewer module is designed to both interactively explore the outputs of Giotto Analyzer and to visualize additional information such as cell morphology and transcript locations (**Fig. 1B**). Giotto Viewer provides an interactive workspace allowing users to easily explore the data in both physical and expression space and identify relationships between different data modalities. Taken together, these two modules provide an integrated toolbox for spatial expression data analysis and visualization.

The spatial omics field is diverse and rapidly expanding; each technology has its strength and weaknesses. In order to demonstrate the general applicability of Giotto, we selected and analysed 10 public datasets obtained from 9 state-of-the-art technologies (**Fig. 1C, Supplementary Table**), which differ in terms of resolution (single-cell vs multiple cells), physical dimension (_2_D vs _3_D), molecular modality (protein vs RNA), number of cells and genes, and tissue of origin. Throughout this paper we use these datasets to highlight the rich set of analysis tools that are supported by Giotto.

### Cell type identification and data visualization

Giotto Analyzer starts by identifying different cell types that are present in a spatial transcriptomic or proteomic dataset. As an illustrating example for the first common steps, we considered the seqFISH+ mouse somatosensory cortex dataset, which profiled 10,000 genes in hundreds of cells at single-cell resolution using super-resolved imaging ^9^. The input gene-by-cell count matrix was first pre-processed through a sequence of steps including normalization, quality control of raw counts, and adjustment for batch effects or technical variations. Then downstream analyses were carried out for highly variable gene selection (**Supplementary Fig. 1B**), dimensionality reduction (such as PCA, tSNE ^19^, and UMAP ^20^), and clustering (such as Louvain ^21^ and Leiden clustering algorithms ^22^) (**Supplementary Fig. 1C**). Cluster-specific marker genes (**Supplementary Fig. 1D-E**) were identified through a number of algorithms (such as Scran ^23^ and MAST ^24^) and a new algorithm based on Gini coefficients ^25^. Whereas the strongest marker genes are typically identified by all three methods, each method has its own strength in detecting specific types of genes (**Supplementary Fig. 2A-G**, see **Methods** and **Supplementary Notes** for details). In total, we identified 12 distinct cell types, including layer-specific excitatory neurons (eNeurons) (*Syt17* in layer 2/3, *Grm2* in layer 4, *Islr2* in layer 5/6), two types of inhibitory neurons (iNeurons) (*Lhx6* vs *Adarb2*), astrocytes (*Gli3*), oligodendrocytes (*Plekhh*), oligodendrocyte precursors (OPCs) (*Sox10*), endothelial cells (*Cldn5*), mural (*Vtn*), and microglia (*Itgam*) cells. The distribution of these cell types can then be visualized in both expression and physical space (**Supplementary Fig. 1F**).

**Figure 2.**
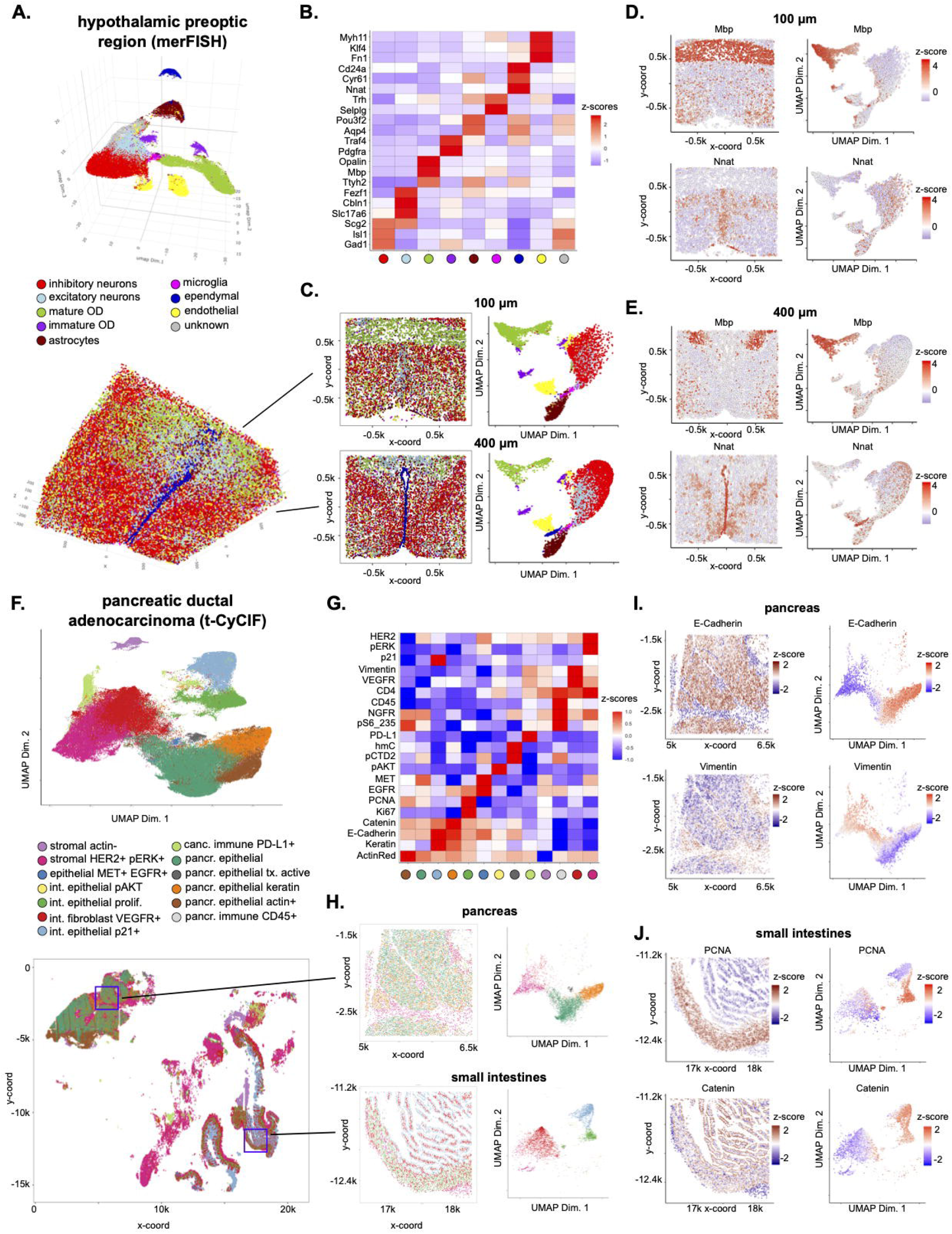
Analysis and visualization of large-scale spatial transcriptomic and proteomic datasets. **A**. Visualization in both expression (top) and physical (bottom) space of the cell types identified by Giotto Analyzer in the pre-optic hypothalamic merFISH dataset, which consists of 12 slices from the same 3D sample (distance unit = 1 µm). **B**. Heatmap showing the marker genes for the identified cell populations in **A. C**. Visualization in both expression and physical space of two representative slices within the z-orientation (100 µm and 400 µm). **D-E**. Overlay of gene expression in both expression and physical space for the selected slices in **C. F**. Visualization in both expression (top) and spatial (bottom) space of the clusters identified by Giotto Analyzer in the pancreatic ductal adenocarcinoma (PDAC) tissue-CyCIF dataset, which covers multiple tissues, including pancreas, small intestine and cancer cells (distance unit = 1 µm). **G**. Heatmap showing the marker proteins for the identified cell clusters in **F. H**. Visualization in both expression and physical space of two selected windows (red squares in **F**.) in the normal pancreas and small-intestinal regions. **I-J**. Overlay of gene expression in both expression and physical space for the selected windows in **H**.

Next, we analysed additional complex imaging-based spatial transcriptomic datasets generated by merFISH ^14^, STARmap ^7^ and osmFISH ^11^. In the merFISH dataset, 12 selected thin slices from a 3D mouse pre-optic cortex sample were imaged, resulting in a total of roughly 75,000 cells and 155 genes. Here Giotto was used to identify 8 distinct clusters. Based on known marker genes, we were able to annotate these clusters as microglia (*Selplg*), ependymal cells (*Cd24a*), astrocytes (*Aqp4*), endothelial cells (*Fn1*), mature (Mbp) and immature (*Pdgfra*) oligodendrocytes, excitatory (*Slc17a6*) and inhibitory (*Gad1*) neurons, respectively, which is in agreement with the original paper ^14^ (**Fig. 2A, B, Supplementary Notes**). Cells that did not fall into these clusters were collectively assigned to an ‘ambiguous’ group, as done in the original paper. Next, to visualize the results Giotto can create an interactive 3D plot for the whole dataset or specifically highlight one or more selected 2D slices (**Fig. 2C**). Together with overlaying gene expression information (**Fig. 2D, E**), such visualization enables the user to explore tissue structure and concomitant gene expression variation in a detailed manner.

In a similar manner we analysed the mouse visual cortex STARmap (**Supplementary Fig. 3 A-D**) and mouse somatosensory cortex osmFISH (**Supplementary Fig. 4 A-D**) datasets (see **Supplementary Notes** for details). Both datasets show the typical anatomical multi-layered structure of the cortex. In the 3D STARmap analysis we present an additional functionality that allows the user to create 2D virtual sections of a 3D sample (**Supplementary Fig. 3 A, C and E**), which could be useful for more refined structural analysis, as demonstrated in our analysis of the STARmap dataset (see section **Giotto identifies distinct cellular neighborhoods and interactions** and **Supplementary Notes** for details).

**Figure 3:**
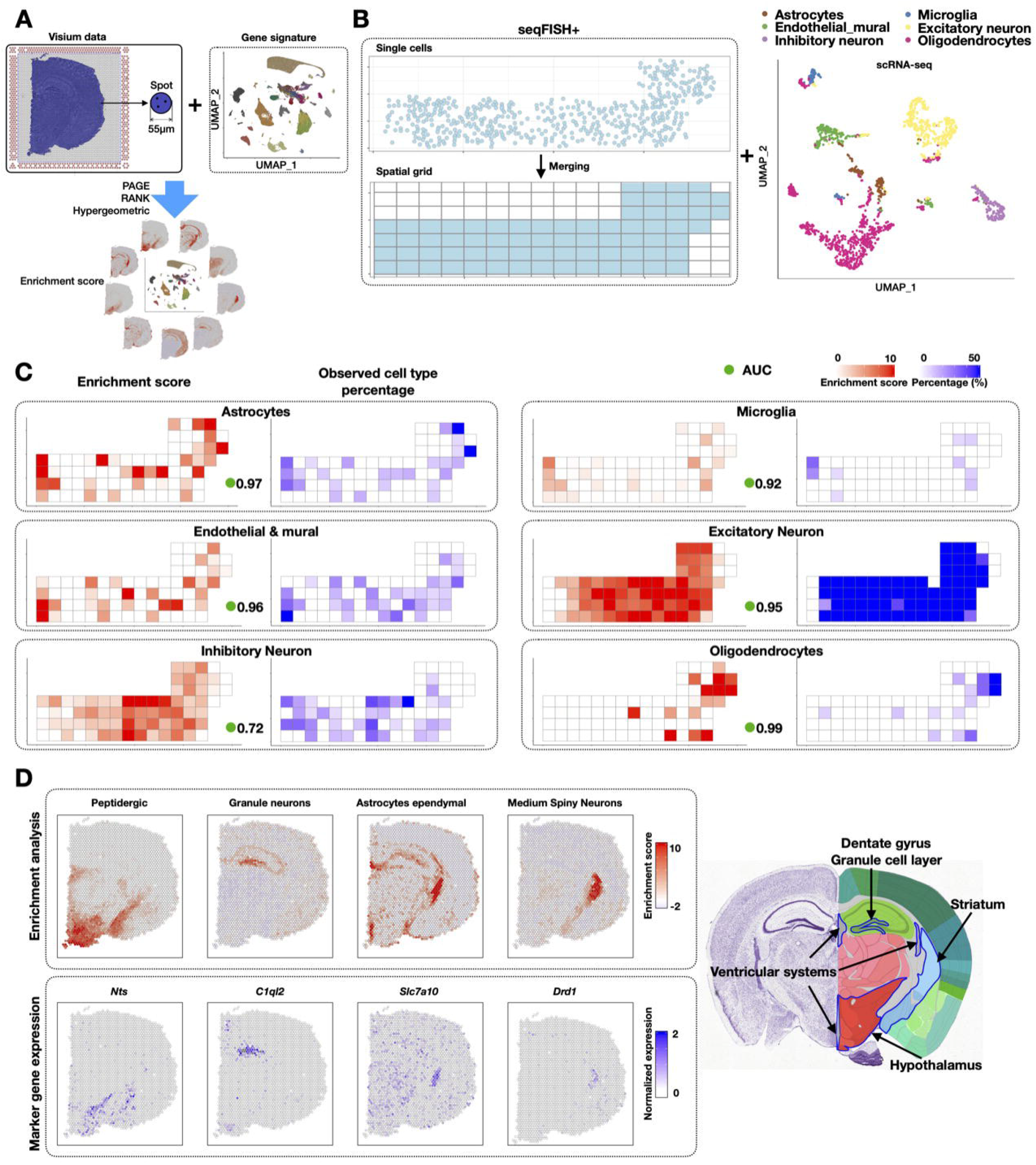
Cell-type enrichment analysis on spatial expression data. **A**. Schematic of cell-type enrichment analysis pipeline. The inputs are spatial expression data and cell type specific gene signatures. These two sources of information are integrated to infer cell type enrichment scores. Giotto implements three methods for enrichment analysis: PAGE, RANK and Hypergeometric. **B**. Single-cell resolution seqFISH+ data are used to simulate coarse-resolution spatial transcriptomic data generated from spot-like squares by projecting onto a regular spatial grid (500 × 500 pixels). Colored squares indicate those that contain cells. External scRNAseq data are visualized by UMAP. **C**. Comparison of cell-type enrichment scores (left, inferred by PAGE) and observed frequency of various cell types (right, based on seqFISH+ data). The agreement between the two is quantified by area under curve (AUC) scores (green circles). **D**. Cell type enrichment analysis for the mouse Visium brain dataset (distance unit = 1 pixel, 1 pixel ≈ 1.46 µm). Enrichment scores for selected cell types are displayed (top left) and compared with the expression level of known marker genes (bottom left). For comparison, a snapshot of the anatomic structure image obtained from mouse Allen Brain Atlas is displayed. Known locations for the selected cell types are highlighted.

**Figure 4:**
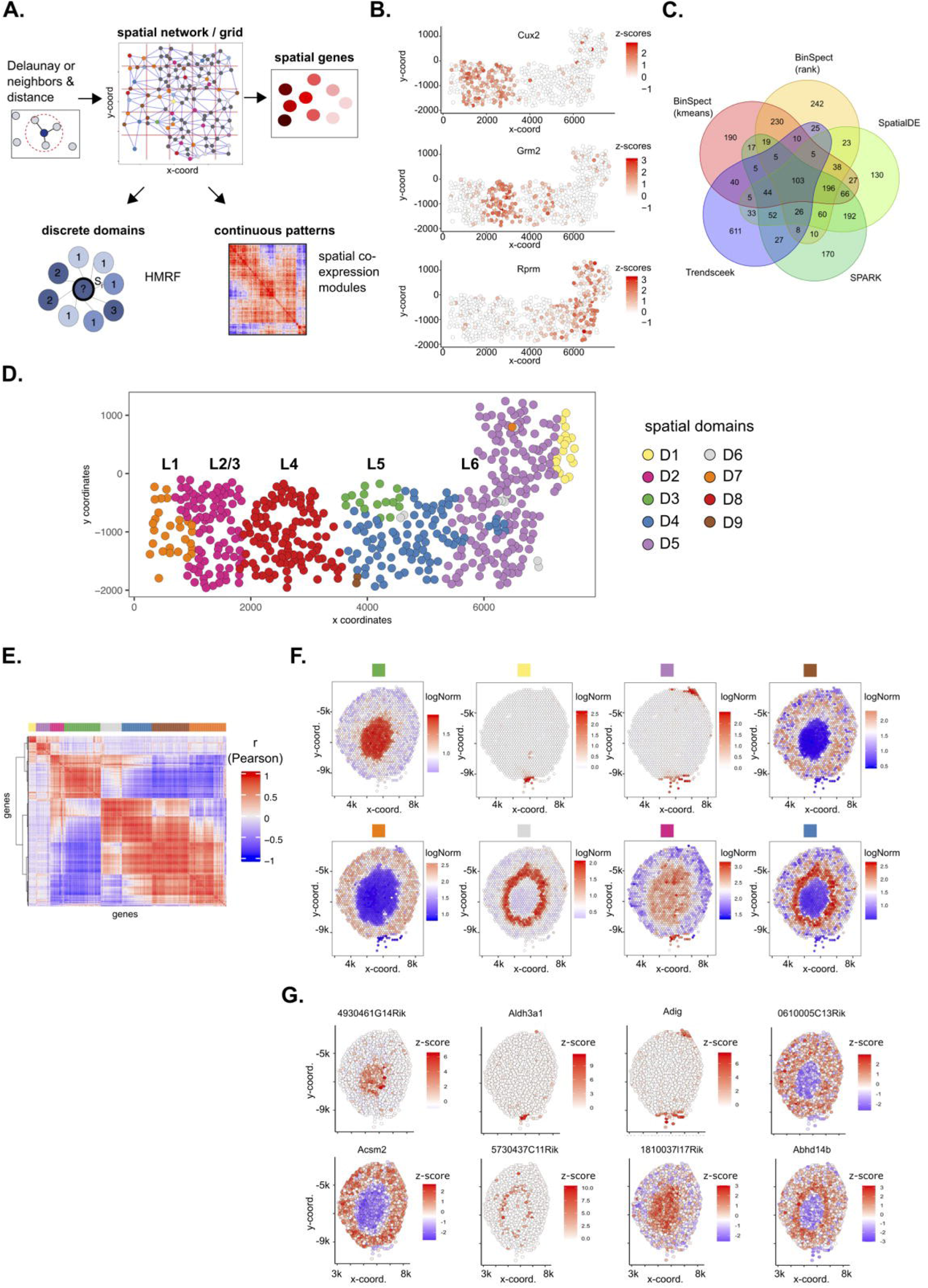
Layers of spatial gene expression variability. **A**. Schematic representation of the subsequent steps needed to dissect the different layers of spatial gene expression variability. The original cell locations, a spatial grid or a spatial network is required to identify individual genes with spatial coherent expression patterns. Those spatial genes can then be used as input to compute continuous spatial co-expression patterns or to find discrete spatial domains with HMRF. **B-D**. Spatial gene expression analysis of the seqFISH+ somatosensory cortex dataset (distance unit = 1 pixel, 1 pixel ≈ 103nm). **B**. Examples of identified spatial genes within the somatosensory multi-layered cortex. The outer layers are on the left, while more inner layers are on the right. **C**. Overlap between the top 1000 spatial genes identified from the 5 methods implemented in Giotto. **D**. Visualization of spatial domains identified by the HMRF model. The layered anatomical structure (L1-6) of the somatosensory cortex is indicated on top. **E-F**. Spatial gene expression analysis of the Visium kidney dataset (distance unit = 1 pixel, 1 pixel ≈ 1.46 µm). **E**. Heatmap showing the spatial gene co-expression results. Identified spatial co-expression modules are indicated with different colors on top. **F**. Metagene visualizations for all the identified spatial gene co-expression modules from **E. G**. Selected gene visualizations for each identified spatial metagene in **E** and **F**.

Due to the similarity of data structure, it is straightforward to apply Giotto to analyse large scale spatial proteomic datasets such as CyCIF, CODEX, and MIBI (see **Supplementary Notes** for details). As an illustrating example, we analysed a public dataset obtained by CyCIF ^10^. The dataset profiled the spatial distribution of 21 proteins and 3 cellular compartment or organelle markers at single-cell resolution in a human pancreatic ductal adenocarcinoma (PDAC) sample that spanned across three distinct tissues: the pancreas, small intestine and tumor. In total, 160,000 cells were profiled. Giotto identified 13 coarse clusters which include mesenchymal, epithelial, immune and cancer cells (**Fig. 2F, G**). Next, we zoomed into each tissue to refine the cell type structure in the pancreas and small intestine separately (**Fig. 2H**). For example, we can now see clearly that the pancreas is structured in distinct zones enriched with epithelial (*E-cadherin*) and mesenchymal or stromal (*Vimentin*) cells, respectively. On the other hand, the small intestine shows a clear proliferating zone (PCNA) and the activation of Wnt signalling (*b-catenin*) in intestinal epithelial cells (**Fig. 2I, J**). Both observations are consistent with the original paper ^10^ Applying the same approach to analyse a mouse spleen dataset from CODEX ^12^ allowed us to identify zones enriched with CD8(+) T cells (Zone 1) and enriched with erythroblasts and F4/80 macrophages (Zone 2) (**Supplementary Fig. 4 E-I**). As such, the employment of Giotto to quickly zoom in to different regions is useful for uncovering the organization of spatial tissues or expression levels in a hierarchical manner.

### Analysis of data with lower spatial resolution

Recently, a number of lower-resolution spatial transcriptomic technologies have been developed, such as 10X Genomics Visium ^26^, Slide-seq ^8^, and DBiT-seq ^27^. Despite their lower spatial resolution, these technologies are useful because they are currently more accessible. To overcome the challenge of lower resolution, Giotto implements a number of algorithms for estimating the enrichment of a cell type in different regions (**Fig. 3A**). In this approach, a continuous value representing the likelihood of the presence of a cell type of interest is assigned to a spatial location which contains multiple cells. To this end, Giotto requires additional input representing the gene signatures of known cell types. Currently, the input gene signatures for the known cell types can either be provided by the user directly as cell type marker gene lists, or be automatically inferred by Giotto based on an additional scRNAseq data matrix input. Giotto then evaluates the match between each cell type’s gene signatures and the expression pattern at each spatial location and reports an enrichment score by using one of the three algorithms: PAGE ^28^, RANK, and Hypergeometric testing (**Fig. 3A**, see **Methods** for details). PAGE calculates an enrichment score based on the fold change of cell type marker genes for each spot. RANK does not require predefined marker genes but instead creates a full ranking of genes ordered by the cell-type specificity score in the scRNAseq data matrix, and computes a ranking-based statistic. Hypergeometric computes the hypergeometric p-value based on the overlap between each cell-type specific marker gene set and the set of spot-specific genes, i.e., those that are expressed at significantly higher levels at certain spots than others (see **Methods**). As negative controls, enrichment scores are also calculated for scrambled spatial transcriptomic data. This allows us to evaluate the statistical significance of an observed enrichment score.

To rigorously evaluate the performance of these cell-type enrichment algorithms, we created a simulated dataset based on the aforementioned seqFISH+ dataset, for which the cell type annotation has been established at the single-cell resolution. To mimic the effect of spatial barcoding, such as that being used in Visium, the merged fields of view were divided into spot-like squares from a regular spatial grid (500 × 500 pixels, ∼51.5 μm) (**Fig. 3B**). For cells located in each square, their gene expression profiles were averaged, thereby creating a new dataset with lower spatial resolution. To apply cell-type enrichment analysis, we obtained scRNAseq data and derived marker gene lists for somatosensory cortex associated cell types from a previous study ^29^. To facilitate cross-platform comparison, we focused on the six major cell-types that were annotated by both studies: astrocytes, microgila, endothelial mural, excitatory neurons, inhibitory neurons, and oligodendrocytes (**Fig. 3B**). For each cell type, we assigned an enrichment score and p-value for each spot by using one of the three enrichment analysis methods mentioned above (**Fig. 3C**). To quantify the performance of each method, we evaluated the area under curve (AUC) score, which was obtained by using the ranking of enrichment score values to predict the presence of a cell type at each spot. Both PAGE and RANK provide high accuracy (median AUC = 0.95 and 0.96, respectively, **Fig. 3C, Supplementary Fig. 5 and 6A**). Even if a spot contains only one cell from a given type, it can often be identified (47 out of 67, ∼70%). The only cell type that cannot be well predicted by this approach is inhibitory neurons, whose gene signatures are less distinct than others.

**Figure 5:**
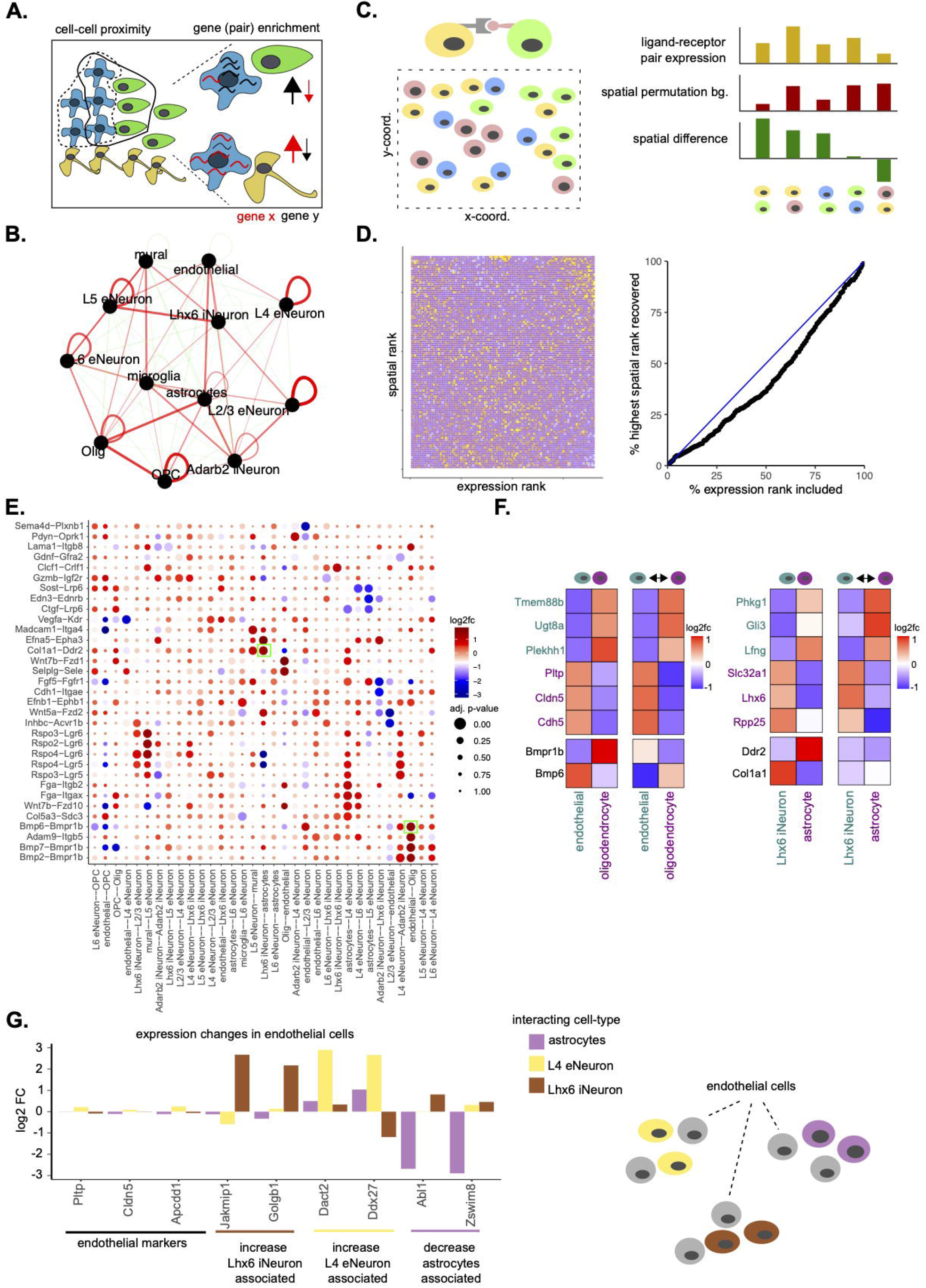
Cell neighborhood and cell-to-cell communication analyses. **A**. Schematic of a multi-cellular tissue with an organized cellular structure (left) and environment specific gene expression (right). **B**. A network representation of the pairwise interacting cell types identified by Giotto in the seqFISH+ somatosensory cortex dataset. Enriched or depleted interactions are depicted in red and green, respectively. Width of the edges indicates the strength of enrichment or depletion. **C**. Visualization of the cell-to-cell communication analysis strategy. For each ligand-receptor pair from a known database a combined co-expression score was calculated for all cells of two interacting cell types (e.g. yellow and blue cells, left). This co-expression score was compared with a background distribution of co-expression scores based on spatial permutations (n = 1000). A cell-cell communication score based on adjusted p-value and log2 fold-change was used to rank a ligand-receptor pair across all identified cells of interacting cell types (right). **D**. Heatmap (left) showing the ranking results for the ligand-receptor analysis as in **C** (y-axis) versus the same analysis but without spatial information (x-axis) for all the ligand-receptor pairs. AUC plot (right) indicating the percentage of expression ranks that need to be considered to recover all the first spatial ranks. **E**. Dotplot for ligand-receptor pairs that exhibit differential cell-cell communication scores due to spatial cell-cell interactions. The size of the dot is correlated with the adjusted p-value and the color indicates increased (red) or decreased (blue) activity. Dots highlighted with a green box are used as examples in **F. F**. Heatmaps showing the increased expression of indicated ligand-receptor pairs between cells of two interacting cell types. **G**. Barplot showing gene expression changes in subsets of endothelial cells (left) stratified based on their spatial interaction with other indicated cell types (right, schematic visualization).

**Figure 6:**
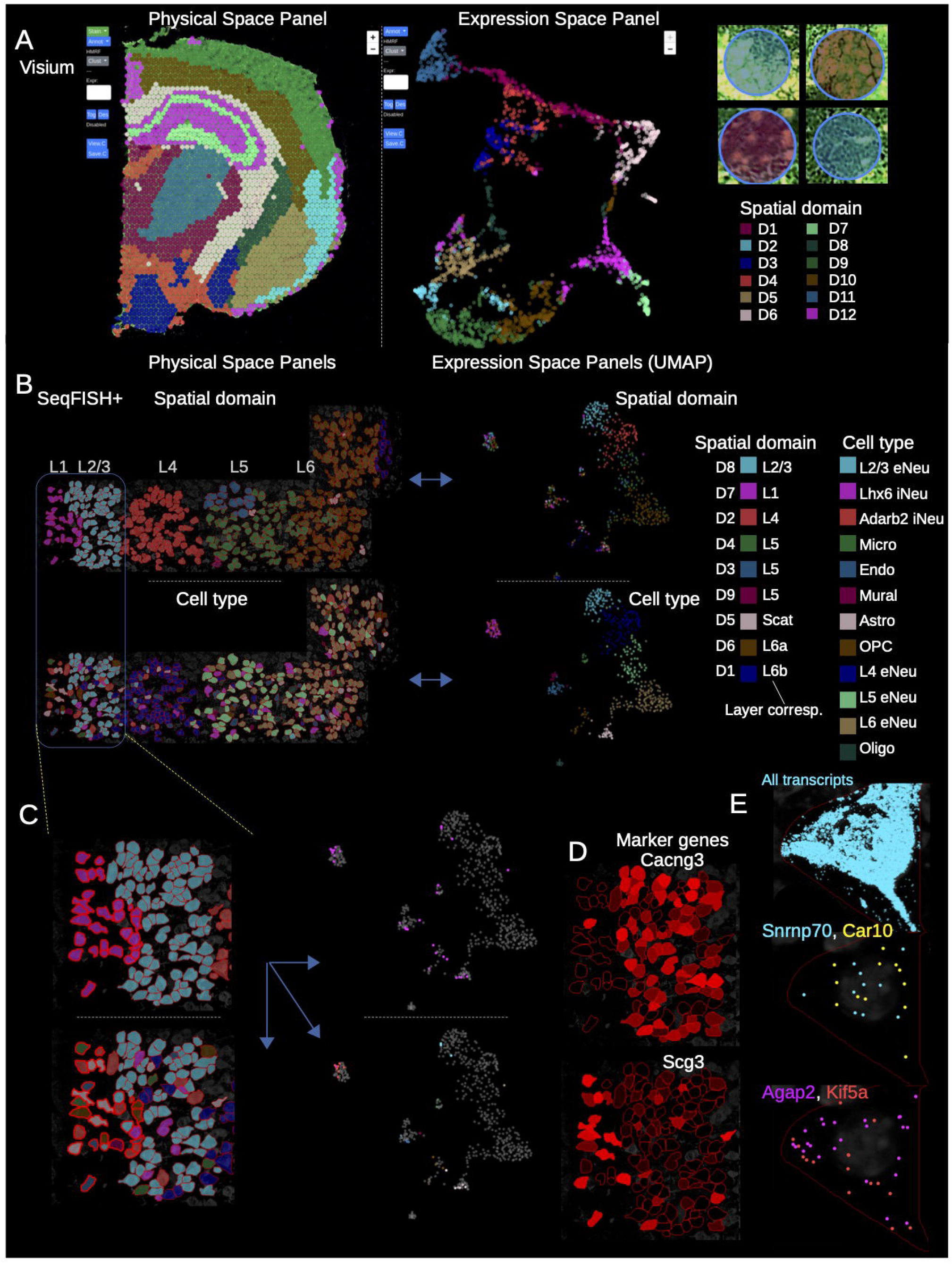
Giotto Viewer provides an interactive workspace to visualize and compare multiple cell annotations. **A**. Visualization of the Visium brain dataset. Two interlinked panels are displayed, showing the data in the physical (left) and expression space (middle). A zoomed-in view shows underlying cell staining pattern at individual spots (right). **B-E**. Visualization of the seqFISH+ mouse somatosensory cortex dataset. **B**. Four interlinked panels are displayed, showing the spatial domain (top) and cell type (bottom) distribution in both physical (left) and expression space (right). **C**. A zoomed-in view of **B**. focusing on the L1-3 regions. Cells in domain D7 are selected (indicated by red outline in left panels and highlighted in the right panels) to enable comparison between spatial domain and cell type annotations. **D**. Expression patterns of representative domain-specific genes. **E**. Subcellular transcript localization patterns of all (top) or selected genes (middle and bottom) in a representative cell. Each dot represents an individual transcript.

In comparison, the hypergeometric and Spearman correlation methods are less accurate (median AUC = 0.86 and 0.72, respectively, **Supplementary Fig. 5** and **6A, Supplementary Notes**). We also compared it with RCTD ^30^, which is a newly developed method for deconvolution. RCTD also performed well (median AUC value of 0.95, **Supplementary Fig 5. and 6A**), but it was considerably slower than the other methods (**Supplementary Fig. 6C**). The four methods that performed well were also robust to changes in number of transcripts (UMIs) (**Supplementary Fig. 6B**).

Next, we applied cell-type enrichment analysis to a publicly available mouse brain Visium dataset (downloaded from https://www.10xgenomics.com/). Spatial transcriptomic information was obtained by using 2,698 spatially barcoded array spots, each covering a circled area with 55μm in diameter. To comprehensively perform enrichment analysis, cell type annotations and corresponding gene signatures were obtained from a public scRNAseq dataset ^31^. Here, we applied PAGE to identify the spatial patterns of the major cell taxonomies identified previously ^31^.

We found that a number of cell types are spatially restricted to distinct anatomical regions (**Fig. 3D, Supplementary Fig. 6D**). The spatial patterns of the enrichment scores are consistent with the literature for a number of cell types, such as peptidergic cells, granule neurons, ependyma astrocytes, and medium spiny neurons (**Fig. 3D**) ^32–34^. Similar but less obvious trends can be observed by inspecting the expression pattern of specific marker genes (**Fig. 3D**), which is consistent with the fact that cell types are typically defined by the concerted activities of multiple genes. Of note, the enrichment analysis also correctly predicted the absence of cell types that should not be present in the sample, such as cerebellum, olfactory bulb, and spinal cord cells (**Supplementary Fig. 6D)**.

To test the general applicability of the enrichment analysis algorithms, we re-analyzed a Slide-seq dataset ^8^ (see **Supplementary Notes** for details), where the read coverage is lower than Visium. This dataset profiles the mouse cerebellum, containing 21,000 beads and 10,500 genes at a coverage of 80 UMIs per bead. Cell-type gene signature information was obtained from a public scRNAseq dataset for a similar region ^31^. Analysis of this scRNAseq dataset identified 15 different cell types. We applied the RANK method for enrichment analysis (**Supplementary Fig. 7A**) and noticed distinct spatial enrichment of cell types in the Slide-seq data that are consistent with prior knowledge. For example, the Purkinje cells were correctly mapped to the Purkinje layer, granule cells were correctly mapped to the nuclear layer, and GABAergic interneurons were mapped to the molecular layer (**Supplementary Fig. 7B**). For comparison, we also applied RCTD to analyze the same dataset and obtained similar results (**Supplementary Fig. 7B, C**).

### Giotto uncovers different layers of spatial expression variability

A key component of Giotto Analyzer is the implementation of a wide range of computational methods to identify spatial patterns based on gene expression. On a fundamental level, Giotto Analyzer represents the spatial relationship among different cells as a spatial grid or network (**Fig. 4A**). To create a spatial grid, each image field is partitioned into regular squares and the gene expression patterns associated with cells within each square are averaged. As such, the spatial grid is a coarse-resolution representation of the data. A spatial network preserves single-cell resolution, and it is created by connecting neighboring cells through a Delaunay triangulation network (see **Methods**). As an alternative approach to create a spatial network, the user can create a spatial network by selecting the k-nearest neighbors or using a fixed distance cut-off, which allows the user to fine-tune the influence of neighboring cells in more downstream applications (**Supplementary Fig. 8A**). However, as shown in the **Supplementary Notes**, our analysis results are typically insensitive to the specific choice of parameter values.

A basic and often first important task in spatial transcriptomic or proteomic analysis is to identify genes whose expression displays a coherent spatial pattern. To this end, Giotto implements a number of methods, including SpatialDE ^35^, Trendsceek ^36^, SPARK ^37^ and two novel methods that are based on spatial network calculations. More specifically the latter two methods are based on statistical enrichment of spatial network neighbors in the high gene expression state after binarization therefore named as BinSpect (**Bin**ary **Sp**atial **E**xtra**ct**ion). The two methods, called BinSpect-kmeans and BinSpect-rank, respectively, differ in the way of binarization (**Supplementary Fig. 8B, Methods**). To evaluate the performance of these methods, we applied each to the seqFISH+ dataset, where many genes are expected to display layer-specific patterns that are consistent with the anatomical structure of the somatosensory cortex (**Fig. 4B, Supplementary Fig. 8C**). For each method, we selected the top 1000 genes as candidates for spatially coherent genes. Of note, a large subset of these genes were identified by at least four of the methods (**Fig. 4C**), these include previously established layer-specific genes such as Cux2, Grm2 and Rprm, indicating that genes with a known and strong spatially coherent expression pattern can be found in a robust manner. On the other hand, subsets of genes were detected by only one or a combination of specific method(s) (**Supplementary Fig. 8D-G**, see **Supplementary Notes**), suggesting it may be beneficial to combine results from all methods for comprehensiveness. The main advantage of the BinSpect methods introduced here is that they are significantly faster compared to SpatialDE (∼ 6-8x), SPARK (∼ 29-45x) and Trendsceek (∼ 816-3300x) (**Supplementary Fig. 8H-I**, see **Supplementary Notes**).

Next, to assess how effective each method could retrieve known spatial patterns we performed a quantitative evaluation based on simulated patterns. In this evaluation we excluded trendsceek since its speed inhibited its use in large-scale simulation studies. Since each method was based on different assumptions or statistical models we established an unbiased simulation strategy based on real high quality data (seqFISH+), which did not rely on a simplified and arbitrary representation of spatial expression data, but instead incorporated all known and unknown variability factors observed in a large subset of genes (**Supplementary Fig. 9**, see **Supplementary Notes** for details). These results show that BinSpect (kmeans and rank) perform systematically better at retrieving known spatial genes, especially when additional noise is introduced. This observation is consistent irrespective of the spatial expression pattern that is evaluated (**Supplementary Fig. 8J)**.

Giotto implements two approaches to systematically summarize the spatial patterns of a large number of spatial genes (**Fig. 4A**). First, Giotto identifies spatial domains with coherent gene expression patterns by implementing our recently developed hidden Markov random field (HMRF) model ^38^. An HMRF model detects spatial domains by comparing the gene expression patterns of each cell with its neighborhood to search for coherent patterns (see **Methods** for details). The inference is based on the joint probability of the intrinsic factor (expression pattern of each cell) and the extrinsic factor (domain state distribution of the surrounding cells) ^38^. The analysis starts with the identification of spatial genes using one of the previously described methods. Then we apply our HMRF model to infer the spatial domain state for each cell or spot. In applying HMRF to the seqFISH+ dataset, our analysis identified 9 distinct spatial domains that were consistent with the anatomic layer structure (**Fig. 4D**). For example, Domain D7 is similar to Layer L1, Domain D2 is similar to Layer L2/3. Of note, such layered structure is not completely reflected by the distribution of different cell types (**Supplementary Fig 1F)**, as numerous cell types (such as inhibitory neurons and endothelial cells) are distributed across multiple layers.

In addition, Giotto also implements a summary view of spatial gene expression patterns based on co-expression analysis. As an illustrating example, we analyzed a Visium dataset obtained from the kidney coronal section, which has known and distinguishable anatomic structures ^39,40^. Using the BinSpect-kmeans algorithm in Giotto, we selected the top 500 spatially coherent genes. To identify spatial patterns, we created a co-expression matrix as follows. First, we spatially smoothed the gene expression data through spatial neighbour averaging, and then created co-expression modules by clustering the spatially smoothed data (**Fig. 4E**). Next, the spatial pattern of each module was summarized by a metagene defined by averaging the expression of all associated genes, which were stored and visualized (**Fig. 4F**). These spatial metagene profiles resemble the known anatomical structures of the mouse kidney and its surrounding environment, which is further corroborated by the spatial co-expressed genes in each module (see **Supplementary Notes**). Moreover, individual genes representing the co-expression patterns were easily extracted and displayed (**Fig. 4G**), providing researchers the opportunity to explore these spatial co-expression patterns in an unbiased manner on a transcriptome wide level. In addition, Giotto also provides a co-expression network based on single-cell expression data so that users can further filter or distinguish spatial co-expression within a local neighborhood from co-expression within the same cell. Finally, these global co-expression patterns ars largely insensitive to the characteristics of the underlying spatial network (**Supplementary Fig. 10A-D**, see **Supplementary Notes** for details).

### Giotto identifies distinct cellular neighborhoods and interactions

Most cells reside within complex tissue structures, where they can communicate with their neighboring cells through specific molecules and signalling pathways. Hence gene expression within each cell is likely driven by the combination of an intrinsic (cell-type specific) component and an extrinsic component mediated by cell-cell communications (**Fig. 5A**). Giotto Analyzer provides a number of tools to explore and extract information related to the cell neighborhood organization, cell-cell communication, and the effect of neighboring cell types on gene expression. To identify distinct cell-type/cell-type interacting patterns, Giotto evaluates the enrichment of the frequency that each pair of cell types are proximal to each other. When analysing the seqFISH+ somatosensory cortex data, we observed that layer-specific neurons usually interact with each other, which agrees with the known multi-layered organization of the cortex (**Fig. 5B**). Such homo-typic (same cell types) relationships are in agreement with what has been observed by others, including in other tissues ^12,41^. Here we also notice that astrocytes and oligodendrocytes, L2/3 and L4 excitatory neurons and L5 and L6 excitatory neurons form frequent hetero-typic (two different cell types) interactions. This is again in line with the expected anatomical structure of the cortex, due to positioning of the cortex layers and the increased presence of astrocytes and oligodendrocytes close to where they originate in the subventrical zone (**Supplementary Fig. 1F** and **Supplementary Fig. 3F**). These observations are robust to changes in number of spatial neighbors (k) (**Supplementary Fig. 11A**, see **Supplementary Notes**) and are furthermore observed in both the seqFISH+ and osmFISH somatosensory cortex datasets (**Supplementary Fig. 11B**, see **Supplementary Notes**).

To extend this type of analysis to less defined tissues, we also analysed a public MIBI dataset profiling the spatial proteomic patterns in triple negative breast cancer (TNBC) patients ^13^. Over 200,000 cells from 41 patients were analysed together to generate over 20 cell populations (**Supplementary Fig. 12A**). Of note, the preferred mode of hetero-typic cell-type interactions is highly patient specific (**Supplementary Fig. 12B, C**). For example, in patients 4 and 5, the Keratin-marked epithelial cells and immune cells are well segregated from each other, whereas patients 10 and 17 feature a rather mixed environment between T cells, Keratin, and Ki67 cancer cells. (**Supplementary Fig. 12B, C**). These observations are consistent with prior findings ^13^.

Giotto builds further on the concept of interacting cell types and aims to identify which known ligand-receptor pairs show increased or decreased co-expression, as a reasonable proxy for activity, in two cell types that spatially interact with each other (**Fig. 5C**). By creating a background distribution through spatially aware permutations (see **Methods**), Giotto can identify which ligand-receptor pairs are potentially more or less active when cells from two cell types are spatially adjacent to each other. By comparing with a spatially unaware permutation method, similar as previously done ^42^, we can see that the predictive power is limited without spatial information (AUC = 0.43) (**Fig. 5C**,**D**). This analysis is relatively stable to different numbers of spatial neighbors (k) within the spatial network (**Supplementary Fig. 11C**, see **Supplementary Notes**) and is observed for multiple ligand-receptor pairs spread out over multiple cell type pairs (**Fig, 5E**). Two potential examples of increased co-expression of a ligand-receptor pair are seen in spatially interacting astrocytes and Lhx6+ inhibitory neurons displaying increased expression of *Ddr2*-*Col1a1* and *Bmp6*-*Bmpr1b* in corresponding endothelial cells and oligodendrocytes, respectively (**Fig. 5F**).

More generally, Giotto implements a number of statistical tests (t.test, limma, Wilcoxon and a spatial permutation test) to identify genes whose expression level variation within a cell-type is significantly associated with an interacting cell type (see **Methods**). After correcting for multiple hypothesis testing, we identified 73 such genes (|log2 FC| > 2 and FDR < 0.1), which we refer to as the interaction changed genes (ICGs). These ICGs are distributed among different interacting cell type pairs (**Supplementary Fig. 11D**). For example, we noticed that endothelial cells interacting with Lhx6 iNeuron were associated with increased expression of *Jakmip1* and *Golgb1*, whereas both *Dact2* and *Ddx27* expression levels were increased in cells from the same cell type but interacting with L4 eNeurons (**Fig. 5G**). On the opposite direction, interaction with astrocytes was associated with decreased expression of *Abl1* and *Zswim8*. Of note, all these subsets of endothelial cells do not show any difference in expression of their known marker genes, such as *Pltp, Cldn5* and *Apcdd1 (***Fig. 5G, Supplementary Fig. 2D**).

### Giotto Viewer: interactive visualization and exploration of spatial transcriptomic data

Giotto Viewer is designed for interactive visualization and exploration of spatial transcriptomic data. Compared to the figure outputs from Giotto Analyzer, the objective of Giotto Viewer is to provide an interactive and user-friendly workspace where the user can easily explore the data and integrate the results from various analyses from Giotto Analyzer, and further incorporate additional information that cannot be easily quantified, such as cell staining images.

Giotto Viewer is a web-based application running in a local environment. It supports a multi-panel view of the spatial expression data. Each panel can be configured to display either the cells in physical or the expression space and overlays gene expression information on top. Complex geometries such as the 2D cell morphology and the associated large antibody staining images of the cells can be toggled easily within each panel. We use a Google Map-like algorithm to facilitate efficient navigation of large data (in terms of either images or cell numbers, see **Methods**). Importantly, panels are interlinked and interactive through sharing of cell ID and annotation information (see **Methods**). This allows seamless integration of different views and facilitates synchronous updates across all panels. We were able to apply Giotto Viewer to display over 500,000 data points (or mRNA transcripts) within a group of cells on one screen without any problem. Indicating the Giotto viewer is capable of handling large datasets.

As an illustrating example, we used Giotto Viewer to visualize the Visium brain dataset. By default, Giotto Viewer creates two panels, representing the data in physical and gene expression space, respectively (**Fig. 6A, left**). Any property that is contained in a giotto object, such as gene expression levels, spatial cell-type enrichment values, cell-type or spatial-domain annotations, can be selected for visualization. Additional imaging-related information, such as cell staining and segmentation, can also be overlaid. The size and location of field of view can be easily adjusted via the zoom and pan functions. At one end of the spectrum, the image content at each single spot can be visualized (**Fig. 6A, right**), revealing the underlying H&E staining pattern. An animated video is provided to illustrate how the user can interactively explore the data and high-level annotations (**Supplementary Video**).

To demonstrate the utility of Giotto Viewer for exploring and integrating a large amount of information generated by Giotto Analyzer, we used the aforementioned seqFISH+ dataset again. Through the analysis described above, we identified various annotations such as cell types, spatially coherent genes, and spatial domains. Therefore, it is of interest to compare the cell type and spatial domain annotation to investigate their relationship. To this end, we created four interlinked panels corresponding to cell type and spatial domain annotations represented in the physical and expression space, respectively (**Fig. 6B**). The view of these panels can be synchronously updated through zoom and pan operations, enabling the user to easily explore the data and inspect any area of interest as desired. For example, as the user zooms in to the L1-L2/3 region (**Fig. 6C**), it becomes apparent that domain D7 consists of a mixture of cell types including astrocytes, microglias, and interneurons. Giotto Viewer also provides a lasso tool that allows users to select cells of interest for further analyses. The borders of the selected cells are highlighted and can be easily traced across different panels. As an example, cells from domain D7 are selected and highlighted (**Fig. 6C)**. By inspecting the pattern in the interlinked panels, it becomes obvious that this domain contains cells from multiple cell types. As such, both cell type and spatial domain differences contribute to cellular heterogeneity.

To gain further insights into the difference between cell type and spatial domain annotations, we saved the selected cells to an output file. The corresponding information was directly loaded into Giotto Analyzer for further analysis. This allowed us to identify a number of additional marker genes, such as *Cacng3* and *Scg3* (**Fig. 6D**). The seamless iteration between data analysis and visualization is a unique strength of Giotto.

In addition, Giotto Viewer also provides the functionality to explore subcellular transcript or protein localization patterns. As an example, we used Giotto Viewer to visualize the exact locations of individual transcripts in selected cells from the seqFISH+ dataset (**Fig. 6E, Supplementary Fig. 12**). To facilitate real-time exploration of the transcript localization data, which is much larger than other data components, we adopted a position-based caching of transcriptomic data (see **Methods**). From the original staining image (**Supplementary Fig. 13A**), the users can zoom in on any specific region or select specific cells and visualize the locations of either all detected transcripts (**Supplementary Fig. 13B**) or selected genes of interest (**Supplementary Fig. 13C**). The spatial extent of all transcripts is useful for cell morphology analysis (**Supplementary Fig. 13B**), whereas the localization pattern of individual genes may provide functional insights into the corresponding genes (**Supplementary Fig. 13C)**. For example, transcripts of *Snrnp70* and *Car10* are preferentially localized to the cell nucleus (delineated by DAPI background), while *Agap2* and *Kif5a* transcripts are distributed closer to the cell periphery (**Supplementary Fig. 13C**).

## Discussion

Single-cell analysis has entered a new phase – from characterizing cellular heterogeneity to interpreting the role of spatial organization. To overcome the challenge for data analysis and visualization, we have developed Giotto as a standardized toolbox, which implements a rich set of algorithms to address the common tasks for spatial transcriptomics/proteomic data analysis, including cell-type enrichment analysis, spatially coherent gene detection, spatial pattern recognition, and cell neighbourhood and interaction analyses. Through analysing diverse public datasets, we have demonstrated that Giotto can be broadly applied in conjunction with a wide range of spatial transcriptomic and proteomic technologies.

Giotto has a number of strengths, including modularization, interactive visualization, reproducibility, robustness, and flexibility. It differs from existing spatial data analysis and/or visualization pipelines ^8,35,36,38,43,44–46^, and is complementary to alternative strategies that computationally infer spatial information from single-cell RNAseq analysis ^47^. To our knowledge, Giotto is the first demonstrated general-purpose toolbox for spatial transcriptomic/proteomic data analysis, while the other methods are designed for specific data types ^44,48–50^ or tasks, such as the identification of cell types ^43^, marker genes ^35,36^ or domain patterns ^38^. Although both Seurat and Scanpy have now adopted spatial transcriptomic analysis functionalities, an important distinction is that our design of the giotto object is specifically targeted for spatial transcriptomic data analysis, whereas the data structure in the other packages were originally designed for single-cell RNAseq analysis. The flexible design of Giotto makes it an ideal platform for incepting new algorithms. In addition, Giotto provides a convenient venue to integrate external information such as single-cell RNAseq data. As single-cell multi-omics data become more available, such integration may greatly enhance mechanistic understanding of the cell-state variation in development and diseases.

## Methods

### Data usage and availability

General information, examples and tutorials are available at http://www.spatialgiotto.com. Users can follow to replicate the analyses in this paper and to learn more about the parameters of specific functions in the Giotto package. Giotto v1.0.2 was used for generating the analyses in this paper. The tutorials give a detailed description of parameters surrounding functions of interest. Dataset examples give a dataset-centric recipe for connecting various analysis components. Download and setup instructions are available. A Docker image has been created and is readily available on http://www.spatialgiotto.com and https://zenodo.org/record/4095230 for ease of access and testing. In addition more detailed information about both the stable and development version of Giotto is available on https://rubd.github.io/Giotto_site/, which is associated with the Giotto source code from our github repository https://github.com/RubD/Giotto. These links also provide additional information about installation issues, documentation, extensive information about how to use Giotto, a news section and guidelines for external contributions.

### Giotto Analyzer

Giotto Analyzer is an open source R package that at its center creates a S4 giotto object (**Supplementary Fig. 1A**), which stores a gene expression matrix, the accompanying cell locations and optionally any associated tissue images for visualization purposes. It contains multiple functions that can either extract or add new information to this object in a flexible manner. In this way users can either follow the default settings or build their own stepwise pipeline to extract and visualize spatial information. In the next part the core steps and functions of Giotto Analyzer are explained and names of functions are depicted in *italic*.

### Quality control, pre-processing and normalization

A giotto object can be created with the function *createGiottoObject*, which requires as minimum input an expression matrix and the spatial coordinates of corresponding cell centroids. If a dataset is composed of multiple fields of view or tiles, they can be stitched together by using *stitchFieldCoordinates* or *stitchTileCoordinates*, respectively. Next, this giotto object can be filtered with *filterGiotto* to exclude low quality cells or lowly expressed or detected genes. As a guide for setting the filtering parameters, the cell and gene distributions can be viewed with *filterDistributions* and the effect of multiple parameters can be tested with *filterCombinations*. Raw counts can further be normalized with *normalizeGiotto*, which can adjust for library size, log transform the matrix and/or perform rescaling. Next the function *addStatistics* computes general cell and gene statistics such as the total number of detected genes and counts per cell. To adjust for variation due to the former technical covariates the *adjustGiottoMatrix* can be applied to the normalized data. If the dataset does not have single-cell resolution, the above steps can still proceed while treating each spatially barcoded spot as a cell.

### Feature selection

To identify informative genes for clustering the *calculateHVG* can be used. Highly variable genes can be detected in two different manners. In the first method, all genes are divided in a predefined number (default = 20) of equal sized bins based on their expression. Within each bin the coefficient of variation for each gene is calculated and these are subsequently converted to z-scores. Genes above a predefined z-score threshold (default = 1.5) are selected for further analysis. For the second method a loess regression model is calculated to predict the coefficient of variation based on the log-normalized expression values. Genes that show higher degree of variability than predicted are considered highly variable genes. Genes can be further filtered based on average expression values or detection percentage, which is returned by the *addStatistics* function.

### Dimensionality reduction

To reduce dimensions of the expression dataset users can perform principal component analysis (PCA) with *runPCA*. Significant PCs can be estimated with the *signPCA* function using a scree plot or the jackstraw method ^51^. Further nonlinear dimension reduction can be performed with uniform manifold approximation and projection (UMAP) ^20^ and t-distributed stochastic neighbour embedding (t-SNE) ^19^ directly on the expression matrix or on the PCA space using *runUMAP* and *runtSNE*, respectively.

### Clustering

First a shared or k nearest neighbour (sNN or kNN) network needs to be constructed with *createNearestNetwork* which uses as input either processed expression values or their projection onto a selected dimension reduction space, such as that obtained from PCA. Louvain and Leiden clustering ^22^ are implemented as *doLouvainCluster* and *doLeidenCluster* and can be directly applied on the created expression-based network. Further subclustering on all or a selected set of clusters can be performed in an analogous manner with *doLouvainSubCluster* or *doLeidenSubCluster*. Alternative clustering options such as kmeans and hierarchical clustering are also available as *doKmeans* and *doHclust*. To aid in further fine-tuning the clustering results users can compute cluster correlation scores with *getClusterSimilarity* and decide to merge clusters with *mergeClusters* based on a user defined correlation threshold and cluster size parameters.

### Marker genes detection

Giotto provides 3 different ways within the function *findMarkers* to identify marker genes for one or more clusters: scran, mast, and gini. The first two methods have been previously published ^24,52^ and are implemented as *findScranMarkers* and *findMastMarkers*. In addition, we also developed a novel method based on the Gini-coefficient ^25^ and implemented it as f*indGiniMarkers*, as schematically illustrated in **Supplementary Figure 2B**. First, we calculate the average log-normalized expression for each gene in each cluster, and represent the result as a matrix *X*, with *X*(*i, j*) representing the average expression of *i*-th gene in *j*-th cluster. Similarly, we calculate the detection fraction of each gene in each cluster, and represent the result as a matrix *Y*, with *Y*(*i, j*) representing the detection fraction of *i*-th gene in *j*-th cluster. For each gene *i*, we calculate two related quantities G_expr_(*i*) and G_det_(*i*), by computing the Gini-coefficient associated with row *X*(*i*, .) and *Y*(*i*, .), respectively. Gini-coefficient of a vector x = [x_1_,x_2_, …, x_n_] is defined as:

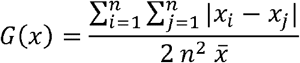

In the meantime, for each gene *i*, we also rank the clusters based on either gene expression *X*(*i*, .) or detection rate *Y*(*i*, .), and the corresponding ranks are denoted by R_expr_(*i*, .) and R_det_(*i*, .), respectively. The ranks are subsequently rescaled between 0.1 and 1. Finally, an aggregated score G_final_(*i, j*) is defined as follows:

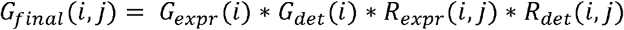

The markers genes associated with a cluster *j* is then identified as those with top values of G_final_(., *j*). We have found this to be a fast and simple approach for effectively identifying genes that are both specific and sufficiently expressed in a particular cluster. In addition, an automated version to perform systematic pairwise comparisons between each cluster and all other clusters is also implemented as *findMarkers_one_vs_all*.

### Enrichment analysis of spatial expression data

Three enrichment methods are implemented into Giotto for enrichment analysis:

#### 1. Enrichment analysis by using PAGE (Parametric Analysis of Gene Set Enrichment) ^28^

In this method, a known set of *m* cell-type specific marker genes is used as input. The objective is to evaluate if these genes are more highly expressed at each spot as compared to other spots. Specifically, for each spot, we define an enrichment score corresponding to a set of marker genes as follows. First, for each gene in the entire genome, we calculate the expression fold change of this gene by using the expression value in this spot versus the mean expression of all spots. Genes associated with high fold change values are annotated as spot-specific genes. The mean and standard deviation of these fold change values are denoted by *µ* and δ, respectively. For comparison, we also calculated the mean fold change associated with the set of marker genes, denoted by *S*_*m*_. The enrichment score (ES) is then defined as follows:

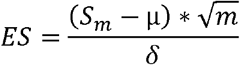

A higher ES value indicates a cell-type is more likely to be associated with the spot in question. To estimate the null distribution, we repeat the analysis by using 1000 random gene sets with the same size. The resulting values are fit by a normal distribution. This null distribution is used to derive p-values associated with the enrichment scores for the real data. The PAGE algorithm is implemented as *runPAGEEnrich*.

#### 2. Enrichment analysis based on rank of gene expression

In this method, a known list of marker genes is not required. Instead, an external single-cell RNAseq dataset is used as input along with the cell-type annotation for each cell. A schematic of this method is illustrated in **Supplementary Figure 7A**. The scRNAseq data matrix is used as input to define cell type specific gene signatures. A fold-change gene ranking is computed for each cell type (scRNAseq) and a relative gene ranking is computed for each spot or bead (Slide-seq is indicated here as an example), which is then used to derive enrichment scores. More precisely, Giotto automatically identifies cell-type specific gene signatures (*makeSignMatrixRank*) by computing the fold change for each gene, *g*, defined as the ratio between its mean expression level within a cell type and the mean level across all cell types, followed by evaluating its relative rank *R1*_*g*_ among all genes. In the meantime, genes are also ranked based on location specificity, *R2*_*g*_, using the spatial expression data. This is obtained by calculating the fold change comparing its expression level at a specific spot versus the overall mean and then ranking the results. The mutual rank ^53^ of gene *g* is then computed as

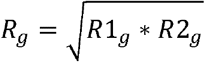

which is then converted to a rank-biased precision (*RBP*) score ^54^, which is defined as follows:

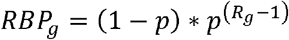

where *p* is a constant set at 0.99. Intuitively, the *RBP* score is used to select genes that are highly ranked in terms of both cell-type specificity and location-specificity. The tuning parameter *p* in the above equation is introduced to control the relative weight of highly ranked genes. In the end, the enrichment score (ES) is determined by the sum of genes with top ranked *RBP* scores:

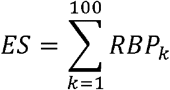

The RANK method is implemented as the *runRankEnrich* function. To estimate the null distribution, we randomly shuffle the ranking of genes in the scRNAseq dataset for 1000 times of each cell type and then apply the above analysis to the shuffled data. The resulting values are fit by a gamma distribution, using the “*fitdistrplus*” package in R. This null distribution is used to derive p-values associated with the enrichment scores for the real data.

#### 3. Enrichment analysis by using hypergeometric distribution

This method also requires a known set of *m* cell-type specific marker genes as input, but it evaluates enrichment by simply using a hypergeometric test. A contingency table is constructed by dividing all genes into four non-overlapping categories, based on marker gene annotation and binarization of gene expression values. The latter is determined by top 5% expression genes for each spot. Based on this contingency table, a *p*-value is calculated. Here the enrichment score is defined as -log10(p-value).

### Simulation analysis of seqFISH+ data

We coarse grain seqFISH+ dataset to simulate spatial expression data of multiple cellular level. The dataset is gridded by 500 pixels by 500 pixels (∼51.5 µm) length (**Figure 3B**). This length is quite similar with the diameter of spots from Visium spatial expression dataset (55 µm). Then the gene expression for each grid is calculated as the sum of normalized single cell expressions within the grid. After normalization of gene expression for coarse-gridded data, we perform enrichment analysis by using somatosensory cortex single-cell RNA-seq data from ^29^ (GSE60361). scRNA-seq data is normalized by *normalizeGiotto* using Giotto. Marker genes for each cell type are identified by using *findMarkers_one_vs_all* with parameter: method = ‘scran’, expression_values = ‘normalized’. We then intersect those marker genes with genes in seqFISH+ data. Top 100 intersect marker genes for each cell type are kept for further enrichment analysis.

To match the cell-type annotations between seqFISH+ and scRNA-seq data, we aggregated the major clusters annotated by Giotto pipeline. ‘L2/3 eNeuron’, ‘L4 eNeuron’, ‘L5 eNeuron’ and ‘L6 eNeuron’ were marked as excitatory neuron (eNeuron). ‘Adarb2 iNeuron’ and ‘Lhx6 iNeuron’ were marked as inhibitory neurons (iNeuron). ‘endothelial’ and ‘mural’ were marked as “endothelial_mural”. The percentage of cell type for each grid was calculated to evaluate the performance of enrichment methods. Top 100 marker genes identified were used for PAGE and hypergeometric analysis by *createSpatialEnrich* with parameters: enrich_method = ‘PAGE’ and enrich_method = ‘hypergeometric’, respectively. The single-cell expression matrix as well as cell labels were used for rank matrix generation by using *makeSignMatrixRank* with default parameters and enrichment analysis by using *createSpatialEnrich* with parameter: enrich_method = ‘rank’.

### Spatial grid and neighborhood network

A spatial grid is defined as a Cartesian coordinate system with defined units of width and height and is created with the function *createSpatialGrid*. The gene expression levels of cells within each grid box are averaged. Another representation of the spatial relationship is the spatial neighborhood network (**Figure 4A**), where each node represents a cell, and each pair of neighboring cells are connected through an edge. The number of neighbors can be defined by setting (a minimal) k and/or radial distance from the centroid position of each cell, and the edge weights can be either binary or continuous. Alternatively, a Delaunay network can be created, which does not require k or radial distance to be specified and is based on Delaunay triangulation. The Delaunay triangulation and its related concept of Voronoi Tessellation was previously applied to study species distribution in the field of eco-geography with the goal to partition a space according to certain neighbourhood relations of a given set of points (e.g. cells) in this space. It has been used and adopted in various fields of biology, including to analyse tissue distribution at the single-cell level ^12^.

### Spatially coherent gene detection

In total Giotto currently has five different methods to identify spatial coherent gene expression patterns. Three previously published methods SpatialDE ^35^, Trendsceek ^36^ and SPARK ^37^ can be run with the functions *spatialDE, trendSceek* and *spark*. The two new methods introduced here are based on statistical enrichment of binarized expression data in neighboring cells within the spatial network, as schematically illustrated in **Supplementary Figure 8B**. First, for each gene, expression values are binarized using kmeans clustering (k = 2) or simple thresholding on rank (default = 30%), which is the only difference between these two methods. Next, a contingency table is calculated based on the binarized expression values between neighboring cells and used as input for a Fisher exact test to obtain an odds-ratio estimate and p-value. In this way a gene is considered a spatial gene if it is usually found to be highly expressed in proximal or neighboring cells. In addition to the odds-ratio and p-value for each gene, the average gene expression, the number of highly expressing cells and the number of hub cells are computed and provided. A hub cell is considered a cell with high expression of a gene of interest and which has multiple high expressing neighboring cells of that gene. These features can be used by the user to further rank and explore spatial genes with different characteristics. We have named the latter method BinSpect (**Bin**ary **Sp**atial **e**xtra**ct**) and implemented it as *binSpect*, within this function the user can choose to use kmeans or threshold ranking to binarize the expression matrix.

### Spatial pattern simulation data

Our strategy for simulating spatial patterns is schematically illustrated in **Supplementary Figure 9**. To generate known spatial patterns based on real spatial data we first randomly select a large number of genes (n = 100) from the seqFISH+ dataset, each with their own unique expression distribution, drop-outs, expression levels and other unknown factors of variation. We then create a fixed spatial pattern, ordered the cells according to their gene expression levels and created two groups. One group (group G) contains all the highest expressing cells and is equal to the size of the spatial pattern and another group (group R) contains all the other remaining cells. Cells within group G are given a probability Pr, while cells in group R are given the probability 1 - Pr. This probability is then used to randomly assign each cell to the fixed pattern, such that with Pr = 1 all the highest expressing cells are being randomly positioned into the spatial pattern, while with Pr = 0.5 this process is completely random. This assignment is repeated multiple times (n = 6) so that a distribution is obtained for a range of probability levels Pr = {0.5, 0.65. 0.8, 0.9, 0.95, 0.99, 1}. Thus, for each pattern we create 4200 simulations (100 genes * 7 probability levels * 6 random samples). This analysis is carried out using the function *runPatternSimulation*.

### Spatial co-expression patterns

To identify robust patterns of co-expressed spatial genes the functions *detectSpatialCorGenes* and *clusterSpatialCorGenes* can be used on the identified individual spatial genes. The first function spatially smooths gene expression through a grid averaging or k-nearest neighbour approach and then calculates the gene-to-gene correlation (default = Pearson) scores. In addition, it also calculates gene-to-gene correlation within the original single-cells to distinguish between spatial and cell intrinsic correlation. The second function performs hierarchical clustering to cluster the gene-to-gene co-expression network into modules and creates metagene scores by averaging all the genes for each identified co-expression module, which can subsequently be viewed using the standard viewing options provided in Giotto.

### Spatial domain detection

Spatial domains are identified with a hidden Markov random field (HMRF) model as previously described ^38^. In brief, HMRF is a graph-based model that infers the state of each cell as the joint probability of the cell’s intrinsic state (inferred from the cell’s own gene expression vector), and the cell’s extrinsic state, which is based on the distribution of the states of the cell’s neighbours. The notion of state is the spatial domain in our case. The neighbourhood graph defines the extent of the neighbour cell influence, together with the parameter beta that defines the strength of the interaction of cells. At the end, HMRF assigns each cell to one of k spatial domains (k to be defined by the user). This HMRF model is implemented in Python and incorporated in Giotto by using the consecutive wrapper functions *doHMRF, viewHMRF* and *addHMRF* to discover, visualize and select HMRF domain annotations respectively.

### Identification of proximal or interacting cell types

To identify cell types that are found to be preferentially located in a spatially proximal manner, as a proxy for potential cell-cell interactions, we use a random permutation (default n = 1000) strategy of the cell type labels within a defined spatial network. First, we label the edges of the spatial network as homo- or hetero-typic, if they connect cells of identical or different annotated cell types, respectively. Then we determine the ratio of observed-over-expected frequencies between two cell types, where the expected frequencies are calculated from the permutations. Associated *p-*values are calculated by observing how often the observed value were higher or lower than the simulated values for respectively increased or decreased frequencies. A wrapper for this analysis is implemented in Giotto Analyzer as *cellProximityEnrichment*.

### Gene expression changes within cellular neighborhood

#### 1. Spatially informed ligand-receptor pairing

To investigate how cells communicate within their microenvironment, Giotto can incorporate known ligand-receptor information from existing databases ^55^. By calculating the increased spatial co-expression of such gene pairs in neighboring cells from two cell types, it estimates which ligand-receptor pairs might be used more (or less) for communication between interacting cells from two cell types. This is implemented in the function *spatCellCellcom*, which is short for spatially informed cell-to-cell communication. More specifically, for each ligand-receptor pair, a cell-cell-communication score S is calculated for every pair of cell types. In particular, for ligand L, receptor R, cell type A, and cell type B, S(L,R,A,B) is defined as the weighted average expression of R and L in all the interacting A and Bs, or in other words in the subset of A and B cells that are proximal to each other (based on spatial network).

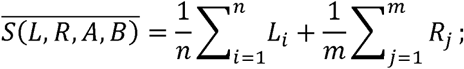

where *n* represents the number of A-type cells that interact with B-type cells, *m* represents the number of B-type cells that interact with A-type cells, Li represents the expression level of the ligand in the *i*-th A-type cell, and Rj represents the expression level of the receptor in the *j*-th B-type cell. Next, to assess if the calculated score *S* is statistically significant, a random null distribution is computed. This background distribution is created by shuffling cell locations within the same cell type for A and B for 1000 (= default) times. In each round a permutation score *Sp* is calculated using the same formula. Associated *p-*values were calculated as the probability of *Sp* to be greater or smaller than the actual observed score *S*. The *p*-values for all ligand receptor pairs in all cell-type pairs were subsequently adjusted for multiple hypothesis testing. A final differential activity score was calculated by multiplying the log2 fold-change with the adjusted *p*-values. The ligand-receptor pair information was retrieved from FANTOM5 ^55^.

#### 2. Expression informed ligand-receptor pairing

This analysis can be performed with *exprCellCellcom*, short for expression informed cell-to-cell communication, and is analogous to the method for ‘spatially informed ligand-receptor pairing’ (described above) except that no spatial information is used. This means that ligand-receptor expression levels are calculated for all cells from two cell types and that the background null distribution is similarly computed by reshuffling all cell labels. This approach is used to mimic scRNAseq based analysis.

#### 3. Comparison between spatially- and expression-information ligand-receptor pairing

We perform a direct comparison between expression-informed ligand-receptor pairing (with the function *exprCellCellcom*) and spatially-informed ligand-receptor pairing (with the function *spatCellCellcom*). For both analyses, we rank all ligand-receptor pairs according to their interaction-induced expression changes in all pairs of cell types. This means that for *spatCellCellcom* we use the cells from two cell types that proximally interact (based on the spatial network) and for *exprCellCellcom* we use all cells from two cell types. Next, we compute the AUC to examine how efficient the expression ranked ligand-receptor pairs can predict the top ranked spatially informed ligand-receptor pair.

#### 4. Cell-type interaction mediated gene expression changes

To identify all potential gene expression changes associated with specific cell-type interactions in an unbiased manner, Giotto implements 4 differential expression tests to identify such interaction changed genes (ICG), including t-test, limma test, Wilcoxon rank sum test, and spatial permutation test. For each cell type, we divide the annotated cells into two complementary subgroups, with one containing the subset which neighbour cells from another specific cell type. Differentially expressed genes between these groups are identified by using each of the statistical tests mentioned above. To adjust for multiple hypothesis testing, a background null distribution is created by reshuffling the cells within the same cell type. This analysis is implemented as the function *findInteractionChangedGenes* or the shorter version *findICG*. Additional filtering can be achieved by using filterInteractionChangedGenes or filterICG in order to reduce errors due to low (interacting) cell number, fold-change or absolute expression differences.

### Giotto Viewer

Giotto Viewer is a web-based application that can be installed on any Linux, Windows or MAC OS based computer. The Giotto Viewer canvas consists of a **cell object layer**, containing all the differently shaped cells, **an image layer** corresponding to staining images, and an **annotation property layer** that specifies the cluster membership and gene expression information. Below is a setup overview:

#### 1. Input files

The minimal input files for Giotto Viewer contain the gene expression matrix and the cell centroid spatial coordinates. Such information can be either provided manually in tabular format, or directly loaded through the output files of Giotto Analyzer. If available, additional input files such as cell segmentations (ROI files), staining images (TIFF files), transcript locations (TXT files), can also be incorporated. Giotto Viewer provides tools to process such information for visualization (see next sections).

To streamline setup, we provide guidelines for setting up various platforms.

i. SeqFISH/MerFISH: this applies to SeqFISH/merFISH where multi-field imaging data, transcript locations, and cell segmentations are available. Giotto Viewer first extracts multi-channel images (where a channel may correspond to Nissl, DAPI, polyA) from TIFF using ImageMagick library (https://imagemagick.org/). Images within the same channel are then stitched. For stitching, Giotto provides an option to stitch images across multiple fields of view (FOV) with gaps in between. The layout can be manually controlled by modifying a coordinate offset file specifying the relative positions of FOVs. To automate these various actions, an initial setup is done using “giotto_setup_image -- require-stitch=y --image=y --image-multi-channel=y --segmentation=y --multi-fov=y -- output-json=step1.json” which creates a template that sets up the sequence of tasks to be performed. Details of the template such as specifying the image file, the stitch offset file, and tiling are next achieved through “giotto_step1_modify_json” script. Lastly, sequence of tasks is performed through “smfish_step1_setup”. An overview of the Giotto Viewer processing steps is in **Supplementary Figure 14**. Cell boundary segmentation is a necessary step for assignment of each transcript to its corresponding cells. However, this task is highly dependent on the specific technology platform therefore not implemented in Giotto. On the other hand, Giotto Viewer can accept user-provided cell boundary segmentation information as input, in the form of Region-of-Interest (ROI) files, for visualization. Giotto Viewer extracts information from the ROI files by adapting a JAVA program based on the ImageJ framework. The next step is tiling the stitched staining image. The purpose of tiling is to support a Google Maps-like algorithm to facilitate multi-level zooming and navigation. Conceptually, this is achieved by tiling of the large image to smaller chunks at various zoom levels to efficiently display a very large image (our stitched image frequently exceeds several hundred megabytes). This is implemented in Giotto Viewer by using the tileup package in Ruby (https://github.com/rktjmp/tileup). This creates a set of tiled images corresponding to 6 zoom levels with 1.5X increment. The size of each tile is fixed at 256 by 256 pixels.
ii. For the Visium platform with an accompanying image, the highest resolution raw H&E staining image is first padded to be square dimension. Next, a template is setup: “giotto_setup_image --require-stitch=n --image=y --image-multi-channel=n -- segmentation=n --multi-fov=n --output-json=step1.json”. The next two steps “giotto_step1_modify_json” and “smfish_step1_setup” are proceeded as usual
iii. Other platforms with no image nor cell segmentations (e.g. Slide-seq). We run “giotto_setup_image --require-stitch=n --image=n --image-multi-channel=n -- segmentation=n --multi-fov=n --output-json=step1.json”, followed by the subsequent two steps (described in ii). Giotto Viewer renders cells as circles in the physical space, and there is not an image background overlay

#### 2. Panels

Giotto Viewer supports a multi-panel view configuration, which means that users can load and visualize any number of panels simultaneously (default number = 2), and add different types of data to each panel. To permit flexibility, there are four types of panels implemented in the Giotto Viewer: *PanelTsne, PanelPhysical*, and *PanelPhysicalSimple, PanelPhysical10X. PanelTsne* requires cell coordinates in the expression space as input. *PanelPhysical* lays out the cells in the physical space in the segmented cell shapes. *PanelPhysicalSimple* is a simplified version of *PanelPhysical* except that cell segmentation and staining images are not required, and instead renders cell objects as fixed size circle markers. *PanelPhysical10X* is unique to Visium in that it also registers the spot-level details of the Visium platform. The number of panels and the panel types can be specified, for example, through: “giotto_setup_viewer --num-panel=2 --input-preprocess-json=step1.json --panel-1=PanelPhysical10X --panel-2=PanelTsne --output-json=step2.json --input-annotation-list=annotation_list.txt”.

#### 3. Annotations

Cell annotations, such as spatial domains and cell types, are required input for Giotto Viewer. In brief, Giotto Viewer supports both continuous- and discrete-value annotations. Annotations generated by Giotto Analyzer can be directly imported by using the *exportGiottoFunction()*.

Once panels, annotations, images are all prepared, then website files are created using: “smfish_read_config -c step2.json -o test.js -p test.html -q test.css”. A python webserver is next launched (python3 -m http.server) and the viewer can be viewed at http://localhost:8000/test.html.

#### 4. Implementation of interactive visualization of multi-layer spatial transcriptomic information

The Giotto Viewer package is written in Javascript utilizing a number of state-of-the-art toolboxes including Leaflet.js (https://leafletjs.com/), Turf.js (https://turfjs.org/), Bootstrap (https://getbootstrap.com/), and jQuery (https://jquery.com/). The Leaflet.js toolbox is used to efficiently visualize and explore multiple layers of information in the data, based on a Google Maps-like algorithm. Leaflet.js recognizes tiles prepared previously by the tileup package and implements caching of tiles and tile handling, allowing it to display large stitched images.

As described above, Giotto Viewer contains 4 types of panels: *PanelTsne, PanelPhysical, and PanelPhysicalSimple, PanelPhysical10X*. The implementation of these 4 panels follows closely the paradigm of object-oriented design in Javascript, specified by the MDN Web Docs and ECMAScript. Briefly, the various panel types are motivated by the fact that depending on data availability, properties of cells change from dataset to dataset, so different ways of cell representation should be considered. In the presence of cell staining images, images should serve as background overlays to the data. If segmentation information is available, cells should be represented in their true cell shape. Yet when neither staining nor segmentation is available, Giotto Viewer represents cells as basic geometric shapes (circles) so that the viewer can still run in the absence of staining or segmentation data. We design the panel classes with these considerations in mind. Giotto Viewer makes it easy for users to specify the number of panels, the type of each panel, and the layout configuration. Users can specify such information in a JSON formatted configuration file. A script then automatically generates the HTML, CSS, JS files of the comparative viewer that is ready for exploration in a standard web browser.

To enable interactivity, panels are linked to each other. This is implemented by first defining mouseover and mouseout events for each cell object. The exact specification of events depends on the type of panel, the action chosen by the user, and the context in which the action is performed. Next, we maintain equivalent cell objects across panels by creating a master look-up table to link cell IDs in different panels. This is useful to facilitate interactive data exploration and comparison, synchronous updates of zoom and view positions during data exploration. Finally, the order and dependency with which interactions are executed are enforced by constantly polling element states and proceeding each step only when states are changed. In the API, the functions *addInteractions(), addTooltips()* enable the easy specification of cross-panel interactions. In the JSON configuration file, interactions between panels are simply defined by the user using the “interact_X: [panel ids]” and “sync_X: [panel ids]” lines.

Giotto Viewer provides an intuitive utility to select a subset of cells of interest for visualization and further analysis. The toggle lasso utility allows a user to hand-draw an enclosed shape in any displayed panel to select cells directly. We implement this function by modifying the Leaflet-lasso.js toolbox to add support for the selection of the dynamic polygon shaped markers. Giotto Viewer can also highlight Individual cells with summary information displayed. This is achieved by using built-in functions in the Turf.js toolbox.

#### 5. Visualizing subcellular transcript localization

To visualize subcellular transcript localization information, an additional layer is created in Leaflet.js. To efficiently handle the large amount of data, we implemented new functions to cache only a small subset of transcripts that fall within the current viewing area, to be rendered by the Leaflet engine and thereby saves the systems resources.

#### 6. Visualizing cell-type enrichment data

Continuous annotations providing cell type enrichment per spot are exported from Giotto Analyzer to Giotto Viewer via the *exportGiotto()* function. Then as with all other annotations, these are added to the setup JSON used to generate the website files. The cell type enrichments will appear under the “Annotations” tab of the viewer panel.

#### 7. Selecting and exporting cells

To encourage iterative analysis between the Giotto Viewer and Giotto Analyzer, users may select any cells of interest with the Lasso button. Then clicking “Save” will save the cell IDs that can be read in Giotto Analyzer within the R environment.

